# First evidence for widespread sharing of stereotyped non-signature whistle types by wild dolphins

**DOI:** 10.1101/2025.04.21.647658

**Authors:** Laela Sayigh, Frants Jensen, Katherine McHugh, Marco Casoli, Laurine Lemercier, Peter Tyack, Randall Wells, Vincent M. Janik

## Abstract

We have built a unique library of sounds produced by known individual common bottlenose dolphins (*Tursiops truncatus*), by recording them non-invasively with suction cup hydrophones during brief catch and release health assessments and with digital acoustic tags (DTAGs). We have catalogued the name-like signature whistles (SWs) of most animals in this resident community of 170 dolphins, which has enabled us to begin studying little known “non-signature whistles” (NSW). We have so far identified 22 shared NSW types, of which two, NSWA and NSWB, are known to have been produced by at least 25 and 35 different dolphins respectively. We are studying the functions of shared NSWs with playback experiments to free-swimming dolphins. We provide background on past playback studies and how they have informed our current research; in particular, received level (RL) of playbacks was found to significantly influence strength of response. Varied responses to playbacks reflect the complexity of dolphin communication, and highlight the need for larger sample sizes to be able to correctly interpret NSW functions. However, results so far have provided support for both the referential nature of SW and the affiliative nature of SW copies (SWCs), because a majority of control playbacks of a dolphin’s own signature whistle (self playbacks) elicited positive responses. NSWA elicited a majority of negative responses, suggesting an alarm-type function, and NSWB elicited varying responses, supporting our suggested function of this whistle type as a “query,” produced when something unexpected or unfamiliar is heard. Given that SW and SWC are known to be learned and appear to be referential signals, it is likely that shared, stereotyped NSW are both learned and referential as well, an idea that is supported by the fact that dolphins are flexible, life-long vocal production learners, unlike most other non-human mammals. Our study provides the first evidence in dolphins for a wider repertoire of shared, context-specific signals, which could form the basis for a language-like communication system.

## Introduction

Bottlenose dolphins, with their large brain-to-body-weight ratios, varied communicative signals, and capacity for lifelong vocal production learning, have long fascinated animal communication researchers. Scientists as far back as the 1960’s engaged in research aimed at discovering dolphin “language” (e.g., Lilly, 1963; Dreher, 1961). However, these studies faced distinct challenges, with the most significant being the difficulty in identifying which dolphin is making a sound – not only are dolphins under water and out of sight much of the time, they also do not make any consistent external movement associated with vocalization. But knowing both who is making a sound and how a receiver responds to it are essential to deciphering any animal communication system.

For the first time in the history of the study of wild cetaceans, we have built a catalog of sounds known to have been produced by specific individual animals. This has been possible due to an innovative research tool employed by the world’s longest-running dolphin conservation research program (Wells, 2009; Wells, 2020): brief catch-and-release health assessments. Since 1984, the Sarasota Dolphin Research Program (SDRP) has been leading these unique research opportunities, during which it has been possible to record known individual common bottlenose dolphins (*Tursiops truncatus*; hereafter referred to as dolphins), identified over decades through photographs of natural markings. This resident community of approximately 170 wild dolphins in the waters in and around Sarasota Bay, FL, USA, spans up to six generations and includes individuals up to 67 years of age (Wells, 2009; Wells, 2020). During health assessments we are able to record dolphins with suction-cup hydrophones placed directly on the melon (Figure 1), resulting in recordings of 313 individuals (57% recorded on multiple occasions) over the past 40 years (Sayigh et al. 2022). Since 2012 we have also been attaching non-invasive suction-cup attached digital acoustic tags (DTAGs, Johnson and Tyack, 2003; Figure 1) prior to release, and now have a obtained more than 100 tag deployments. These powerful data sets have been used to study individually distinctive signature whistles (SW), which are similar to human names (Janik & Sayigh, 2013); previous experimental studies have demonstrated a referential use of such signals (Richards et al., 1984; Harley, 2008). We now have a catalog of SW of most of the dolphins living in this resident community (in addition to many who are no longer living; Figure 2), and much has been learned about these whistles in the past several decades (reviewed in Sayigh et al. 2022). For example, we know that dolphins can copy each other’s SWs as a mechanism to initiate contact (King & Janik, 2013; King et al. 2013), providing additional support for referentiality. We also know that female dolphins increase the maximum frequencies of their SW when communicating with their calves, similar to human “motherese,” or child-directed communication (Sayigh et al. 2023).

**Figure 1:**
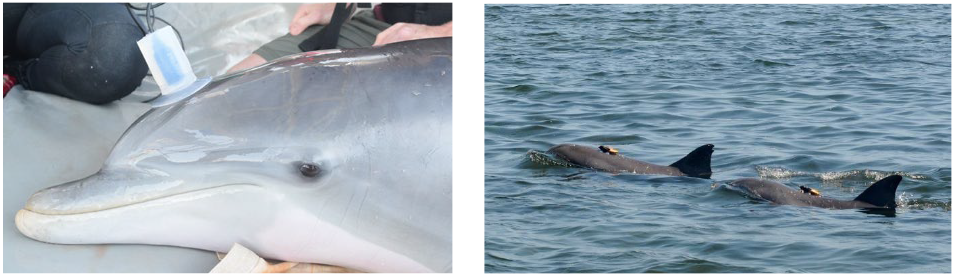
Non-invasive methods for obtaining recordings from known individual wild dolphins. Left: Common bottlenose dolphin being recorded with a suction-cup hydrophone directly on the melon, while being temporarily held during health assessments in Sarasota Bay, Florida; Right: Dolphins wearing suction-cup attached digital acoustic tags (DTAGs). Photos taken by Brookfield Zoo Chicago’s Sarasota Dolphin Research Program under NOAA/National Marine Fisheries Service Scientific Research Permit

**Figure 2:**
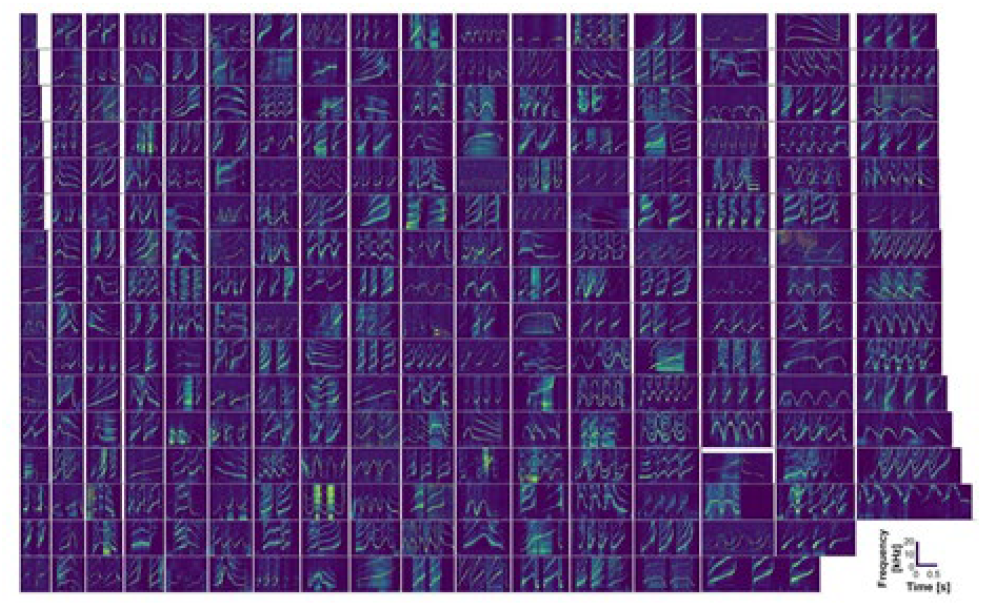
A single example spectrogram of a signature whistle for each of 269 individual bottlenose dolphins, recorded with suction-cup hydrophones in Sarasota, Florida. Time and frequency axes are standardized

Several studies of free-swimming dolphins in Sarasota have estimated the percentage of all whistles that are SW to range between 34 and 70% (Buckstaff, 2004; Cook et al. 2004; Watwood et al. 2005), averaging roughly 50%. Thus, around 50% of whistles produced by these dolphins are “non-signature whistles.” Non-signature whistles (NSWs) have been historically difficult to study because they can only be identified if the SWs in a population are known. But because the SWs of most animals in the Sarasota Bay dolphin community are known, we are uniquely positioned to study NSW.

Before going into more detail about our ongoing work with NSW, we will first describe results from previous (both published and unpublished) playback experiments with dolphins in Sarasota over the past several decades, to illustrate the power of these non-invasive methods to provide insights into dolphin communication, and to provide background for our current work.

### Recording and playback methods

During catch-and-release health assessments, a 500 x 4 m net is deployed from a small outboard vessel in shallow (<2m) water, creating a net corral that contains a small group (1–4) of dolphins for short (1–4 h) periods of time, during which they are continuously monitored by veterinarians. As noted earlier, we record dolphins with non-invasive suction-cup hydrophones (High Tech, Inc.; recorded at 96 kHz sample rate) throughout this period (Figure 1). We selected exemplars from these recordings to use as playback stimuli in our experiments, as well as from recordings made with DTAGs (Figure 1). These multi-sensor archival tags record sound (240 kHz sample rate), depth, and movement, with a magnetometer and 3-axis accelerometers to measure the animal’s orientation (pitch and roll; Johnson and Tyack, 2003).

Natural whistle playback stimuli over the years have consisted of SW, signature whistle copies (SWC), NSW, and unfamiliar whistles. The number of stimuli played back has varied in different experiments, as described in more detail below. Stimuli were high-pass filtered at 2-4 kHz to remove low frequency noise and downsampled to 44.1kHz. For most experiments, a LL9162 underwater speaker (Lubell Labs, Columbus, OH) connected to a power amplifier was used to play back sounds to the dolphins, with sound files played from a netbook or laptop computer. Frequency response for the combined system was 240–20,000 Hz ±3 dB.

We have carried out a series of playback experiments to temporarily-held dolphins during brief catch-and-release health assessments that have provided extensive data on the structure and function of dolphin SWs, and have opened doors to new research questions. All of these experiments followed a similar protocol, in which two 30-sec sequences of whistles were played back, each followed by 5 minutes of silence. The timing of whistles in these playback sequences was modelled after natural signature whistle bouts (Janik et al. 2013). The subject, or ‘target animal’, would be held alongside the boat, so that turning responses toward the speaker could be observed and filmed.

Using this protocol, Sayigh et al. (1999) played back SWs of relatives and familiar associates, and found that dolphins responded more strongly to whistles of relatives. Because familiarity could be ruled out as the driver of these responses, these results were interpreted as evidence for SWs functioning in individual recognition. We next explored the features of whistles that dolphins used for recognition. We aimed to test whether dolphins use “voice cues”, as do many terrestrial mammals (e.g., Boughman & Moss, 2003), to recognize other individuals. Janik et al. (2006) replicated the methodology used by Sayigh et al (1999), but played back synthetic SWs with all potential voice cues removed instead of natural SW. They found the same significant difference in response to synthetic SWs of relatives and familiar associates, showing that dolphins could recognize SWs by means of the distinctive pattern of frequency modulation, or contour, alone. Finding that dolphins were capable of recognizing whistles based on contour alone did not rule out the possibility that they were also capable of utilizing voice cues for recognition, so we explored this by carrying out playbacks of NSWs. Again using the same protocol as Sayigh et al. (1999), Sayigh et al. (2017) played back variable NSW types, and presumed that if dolphins could discriminate whether these NSWs were produced by relatives or close associates, they must be using voice cues to do so, because we did not find any consistent contour cues that would indicate kinship or association. These experiments found that, unlike when hearing playbacks of signature whistles, dolphins did not respond differently to NSW of relatives vs. familiar but unrelated individuals, and thus Sayigh et al. (2017) concluded that dolphins were not using voice cues in individual recognition.

Although we did not find evidence that dolphins recognized the identity of the vocalizing dolphin in our NSW playbacks, their responses to these playbacks opened doors to unexpected new research directions. First, Sayigh et al (2017) found that dolphins often copied NSW playback stimuli, and in one case even engaged in a call-and-response type exchange with an unusually noisy stimulus (Figure 3). We have begun to review past experiments to determine how common this type of behavior is in dolphins (our earlier studies focused on physical turning responses rather than vocal responses to playbacks). Preliminary results indicate that it occurs regularly; for example, a recent playback to a juvenile male dolphin of his mother’s SWs resulted in a prolonged (15 minute) call-and-response exchange in which the juvenile responded to all 148 exemplars of his mother’s SW with his own SW (with response defined as whistling within 1 sec of the end of a playback stimulus; Nakahara & Miyazaki, 2011; Figure 4). These call interactions are not just an automatic response; during the exchange we recorded the same animal interacting vocally with a more distant conspecific (he produced one of the shared NSW types described in more detail below). We plan to further document both NSW copying and call-and-response behaviors in more detail in future work.

**Figure 3:**
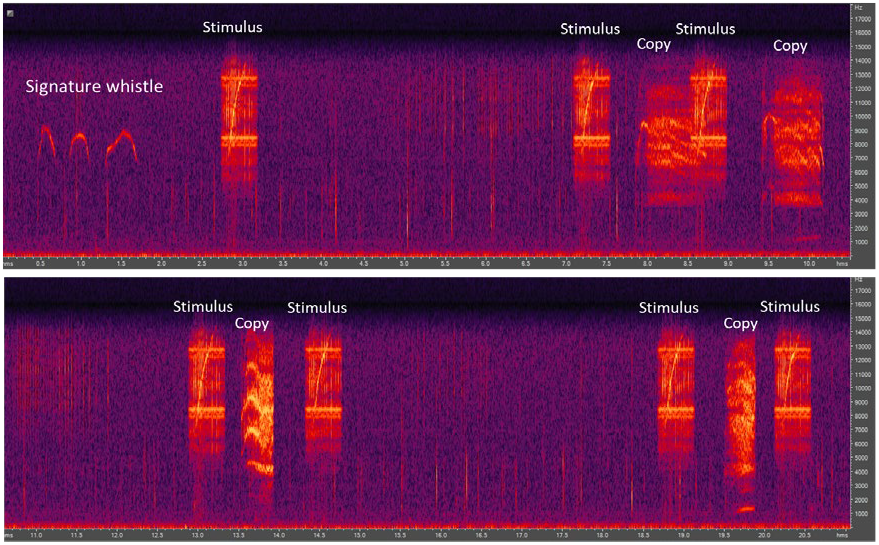
Spectrogram of copying of a noisy non-signature whistle playback stimulus; a 22-s sequence is divided into two 11-s sections. The target animal’s signature whistle is visible at the beginning, followed by a stimulus presentation and then several stimulus-copy exchanges. Frequency (up to 18,000 Hz) is on the y axes, and time in seconds is on the x axes.

**Figure 4:**
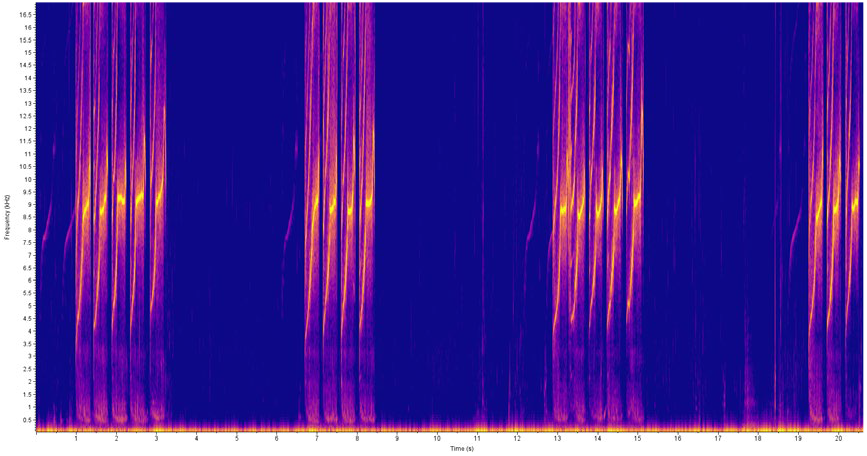
Spectrogram of a 20-sec excerpt of a 15 minute whistle exchange between a juvenile dolphin (louder whistles) and playback stimuli of his mother’s signature whistle (quieter whistles). Frequency (up to 17,000 Hz) is on the y axis, and time in seconds is on the x axis

Another intriguing finding was our first observation of a shared, repeated NSW type, which occurred in response to playbacks of NSWs (Sayigh et al. 2017). Six male dolphins were found to produce a similar NSW (NSWB) in response to these playbacks; this whistle type is characterized by a variable beginning portion followed by a prominent flat (constant frequency) portion, typically centered around 3kHz (Figure 5). Constant frequency whistles are unusual in Sarasota (Miksis et al. 2002) so these whistles stood out as responses in these experiments. Since then, we found additional evidence for NSWB (more details below), and it was this finding that led to our current systematic effort to build a catalog of stereotyped NSWs that are shared by more than one individual.

**Figure 5:**
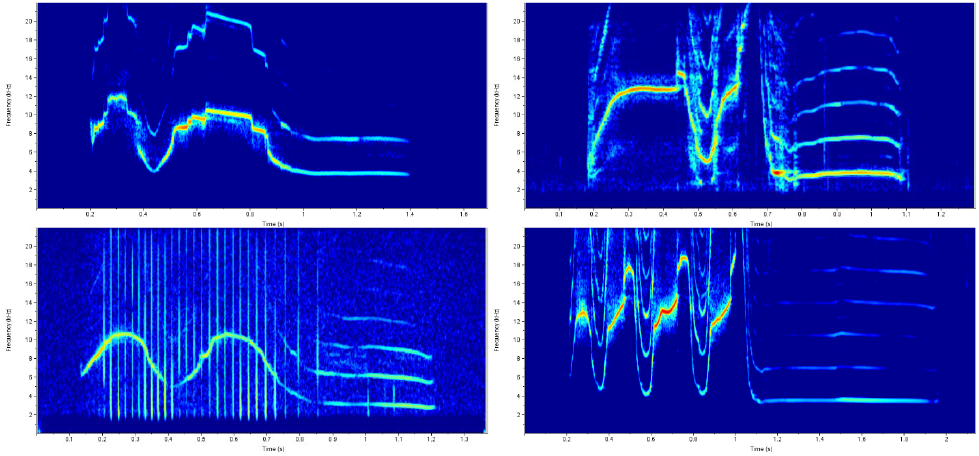
Four examples (produced by four different dolphins) of NSWB, characterized by a variable beginning followed by a constant frequency portion, typically at around 3kHz. Frequency (up to 22,000 Hz) is on the y axes; time in seconds is on the x axes

### Identification of repeated NSW types

Whistles recorded during health assessments were labelled as either SW or NSW (in addition to several other categories including SW copy (SWC) and SW variant) and extracted into a database (see methods in Sayigh et al. 2022). For our current work, we focused on whistles produced by a subset of 117 animals still living in the community for which we had health assessment recordings within the past 20 years, with the goal of selecting whistle exemplars to use in playback experiments. High quality NSWs were reviewed, and repeated contours were visually classified according to contour shape (visual classification of whistles has been found to be highly accurate; Janik, 1999; Sayigh et al. 2007). A similar process was used to identify repeated NSW in DTAG recordings.

To date, we have identified 22 shared NSW types that are produced by three or more individuals. NSWB (Figure 5) has so far been found to be produced by at least 35 different individual dolphins, and NSWA (Figure 6) by at least 25. Interestingly, several of these NSW types are unlike any SW in our extensive SW catalog (Figure 2), and are characterized by highly unusual features. As previously mentioned, the constant frequency portion that characterizes NSWB is a feature rarely seen in SWs of wild dolphins (Figure 6; Miksis et al. 2002). The short and steep up- and down-sweeps that characterize the structure of NSWA (Figure 6) are also uncommon in SW. NSWC (Figure 7) contains both of these unusual features, consisting of short, steep up- and down-sweeps on either side of high, relatively constant frequency components. Although not all shared NSW types contain unusual features like these, we aim to explore whether these features may be related to how these whistles function. Figure 8 shows 10 examples of NSW types with more “typical” (SW-like) contours.

**Figure 6:**
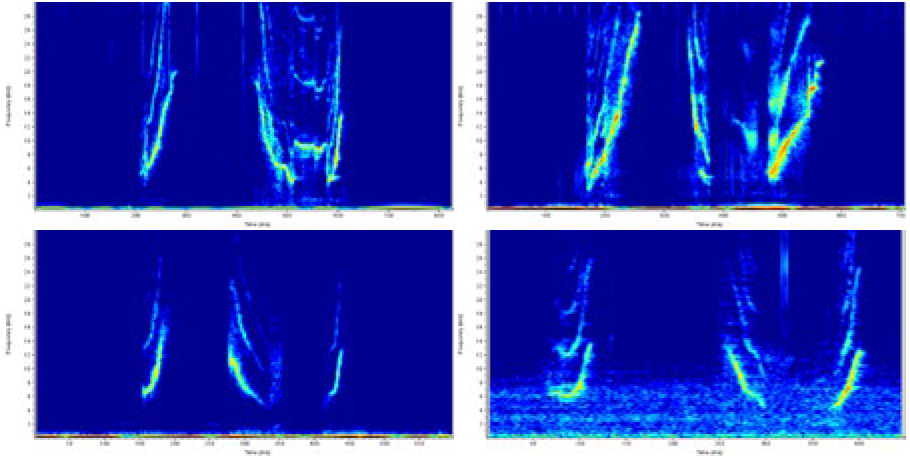
Four examples (produced by four different dolphins) of NSWA, characterized by a steep upsweep, a steep downsweep, and another steep upsweep. Frequency (up to 22,000 Hz) is on the y axes; time in seconds is on the x axes

**Figure 7:**
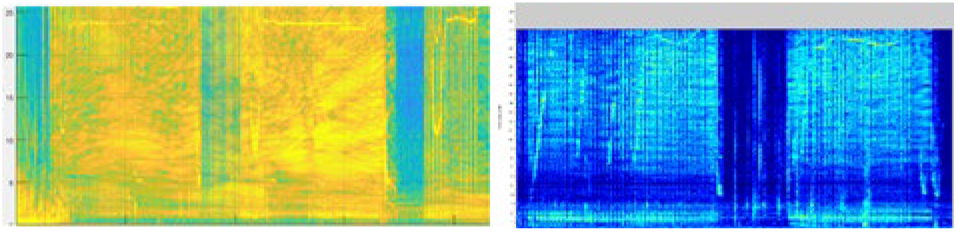
Two examples of NSWC, which consists of steep up- and down-sweeps separated by relatively flat, high frequency segments; these whistles typically co-occur with echolocation clicks (which appear as yellow vertical lines or smears). Whistle on the left was recorded on a DTAG, whereas the example on the right was recorded at a moored hydrophone in Sarasota Bay (this hydrophone only recorded up to 22kHz, but the scale was adjusted to be the same as the spectrogram on the left (up to 25kHz), for comparison purposes); each whistle is approximately 3 seconds in duration

**Figure 8:**
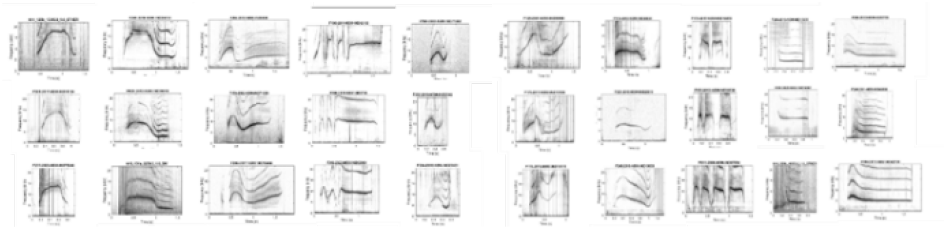
Ten examples of shared NSW types; each column is a different type, with the exemplars on each row produced by different dolphins

### Playbacks to free-swimming dolphins

Playback experiments that were described above were carried out under controlled circumstances during brief catch-and-release health assessments. However, to try to unravel how these shared NSW types function in the dolphin communication system, it is necessary to play back sounds to free-swimming dolphins, to be able to observe their full range of possible responses. Some earlier playback trials with free-swimming dolphins that focused on other research questions were critical in informing our NSW playback protocol, so some aspects of these trials will be described, after first describing the basic playback protocol and methodology.

Once a free-swimming candidate for playbacks was identified (either during boat-based surveys, or when following animals wearing DTAGs after release), we would stay with the animal for approximately 30 minutes, including a 10-minute pre-trial period during which systematic behavioral observations were recorded at 3 minute intervals. After this pre-trial period, a drone (usually a DJI phantom 4 Pro V2.0) would be launched, and a hydrophone deployed (High Tech, Inc., recorded with 96 kHz sample rate). When the target dolphin was positioned approximately 50-100m away and approximately abeam of the boat, the boat was taken out of gear and the speaker (Lubell labs LL9162) suspended at approximately 1 m depth for playback. Playback sequences consisted of two repetitions of the same whistle separated by 2-3 seconds of silence (which is a typical inter-whistle interval; Janik et al. 2013). We aimed to expose each dolphin to two trials, separated by at least 15 minutes, although in some cases we took advantage of an animal wearing a DTAG to carry out one or two additional trials, for a maximum of four trials per dolphin. Continuous behavioral observations were carried out throughout all trials and inter-trial intervals, and for at least 10 minutes following the last trial.

Our first playback trials to free-swimming dolphins focused on male alliances, which in Sarasota are closely bonded pairs of males (Owen et al. 2002). The goal of these initial experiments was to examine how members of an alliance, when voluntarily separated from each other, would respond to playbacks of their alliance partner’s SW, and of their alliance partner copying their own SW (Casoli 2023). In some cases we would also do a third trial that consisted of whistles from a completely unfamiliar individual. Although it was challenging to obtain a large enough sample size for these experiments, due to the rarity of male alliances from which we had recorded SWC separating from each other, these experiments still provided valuable insights into our playback protocol, which will be described in more detail below in the section on playback amplitude levels. In addition, these playbacks provided a fascinating example of NSW production that is worth describing even though it occurred only two times in a pair of playback trials with a single male alliance. In this instance, the male alliance was staying close together, but we went ahead with the playback, in order to practice our protocols (drone launch, speaker deployment, etc.). These two males were close together when they heard first a playback of one of their own SW, and 15 minutes later a playback of one of them copying the other’s SW. In both instances we recorded a faint but detectable rendition of NSWB as a response (Figure 9), with the characteristic low (appr. 3kHz) and constant frequency portion at the end of the whistle. In fact, there appeared to be overlapping contours at 3kHz, suggesting that possibly both males made a similar NSW response. Although anecdotal, this finding provided further impetus to pursue targeted NSW playback trials.

**Figure 9:**
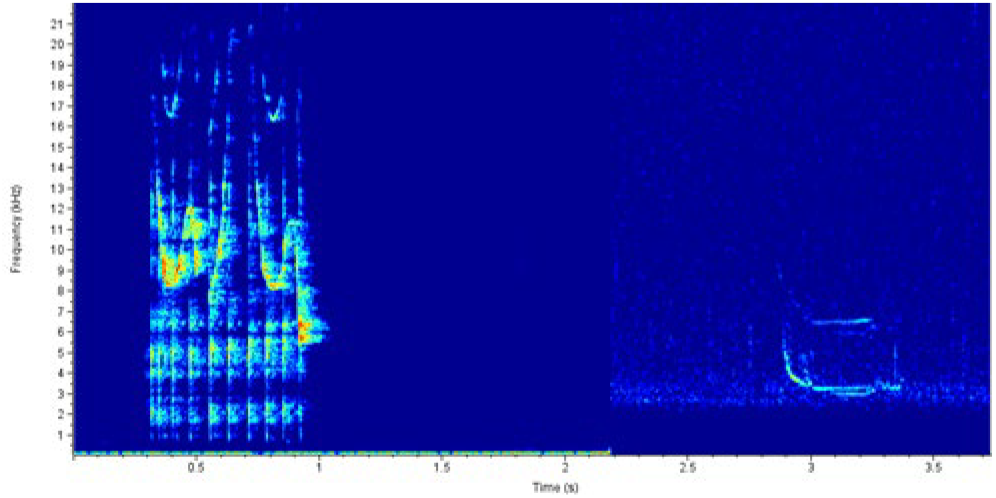
Spectrogram of a playback of the signature whistle of one member of a male alliance, which was played back to him and his partner while they were together. Approximately 2 sec following the playback, a faint but detectable whistle that contains features of NSWB (flat portion at the end at approximately 3kHz) was recorded (note: this section was high-pass filtered and amplified to increase resolution). Although difficult to discern, it is possible that both males are simultaneously producing a similar NSW response, as evidenced by overlapping contours

### Playback source and received level

As noted above, these early trials provided key insights into our playback protocol, especially surrounding the appropriate source level for our stimuli. We initially utilized a source level of 150dB re 1 µPa RMS, which was used for whistle playbacks to free swimming dolphins by King and Janik (2013), and is within the range of whistle source levels measured on DTAGs (Kragh et al. 2019). We controlled stimulus source levels by recording speaker output with a calibrated Soundtrap recorder (Ocean Instruments, NZ) prior to the start of experiments, to determine the system settings needed to achieve our desired source level. Stimuli were normalized to this pre-determined level in Adobe Audition software.

In early trials utilizing 150dB re 1 µPa source levels we often found surprisingly strong avoidance responses, even when playing back familiar whistles. We considered the possibility that dolphins are likely aware of which other animals are in their vicinity, so when they hear another animal apparently nearby so suddenly it could be surprising and disorienting. So in subsequent trials we began playing whistles at reduced source levels of 135-140 dB re 1 µPa. Focusing on a subset of trials that were carried out with tagged dolphins (47 experiments on 20 different dolphins), where the stimulus received levels (RLs) could be measured directly from tag recordings, we found that RLs fell into two significantly different categories (t test, p<0.001): greater than 120dB re 1 µPa and less than 120dB re 1 µPa (Figure 10), corresponding to experiments carried out with 150dB re 1 µPa vs. lower source levels. As a response category, we examined Overall Dynamic Body Acceleration (ODBA), which can be calculated from the accelerometers on the tags, and can be used as a proxy for animal movement. Stimuli were divided into two broad categories: familiar (self, SWC, familiar SW) and unfamiliar (whistles from a different population)^1^. ODBA was found to significantly increase following playback across all trials (Wilcoxon signed-rank test for paired data, p<0.001; Fig 11); however, ODBA was significantly higher in response to stimuli in the higher vs. the lower RL category, regardless of stimulus type (Kruskal-Wallis rank sum test p=0.02; Fig 12). Thus, this analysis was critical in our shift to lower source level playbacks when we began our NSW trials.

**Figure 10:**
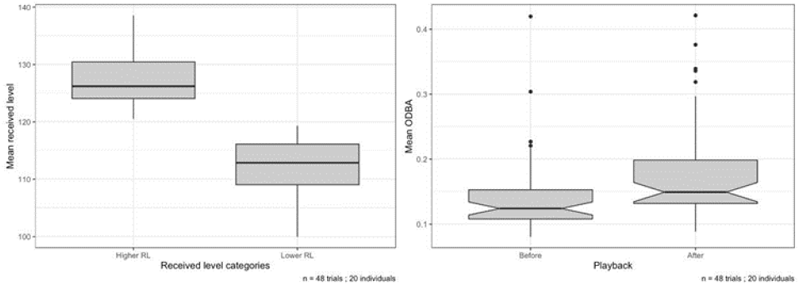
Left: Mean received level (RL) as measured from tag recordings for 48 trials on 20 individual dolphins. RLs fell into two significantly different categories: greater than 120dB and less than 120dB (t test, t=10.2, p<0.001); Right: Overall Dynamic Body Acceleration (ODBA) was significantly higher following playbacks than before (Wilcoxon signed-rank test for paired data, p<0.001)

**Figure 11:**
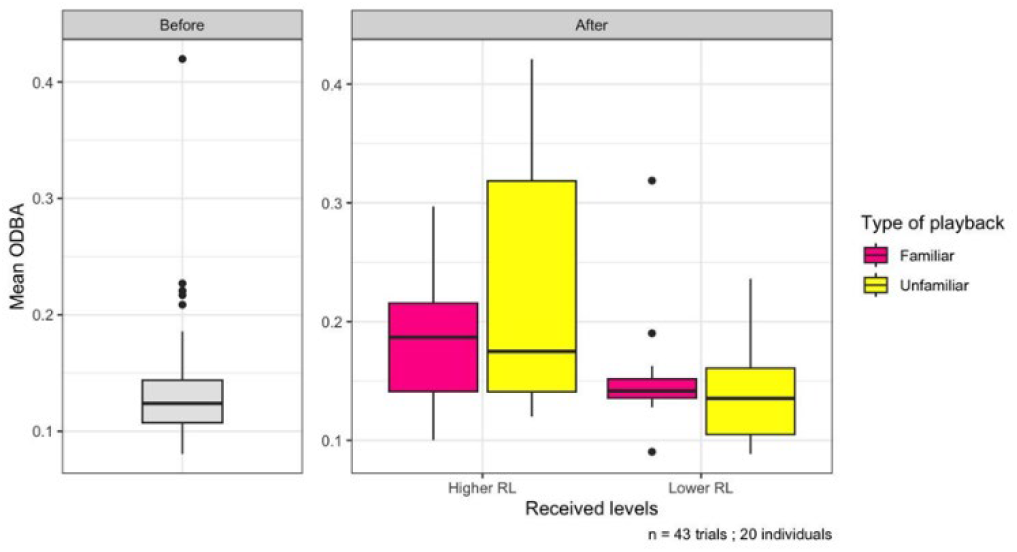
Mean Overall Dynamic Body Acceleration (ODBA) was significantly higher following higher RL playbacks, regardless of type (Kruskal-Wallis rank sum test, p=0.02)

**Figure 12.**
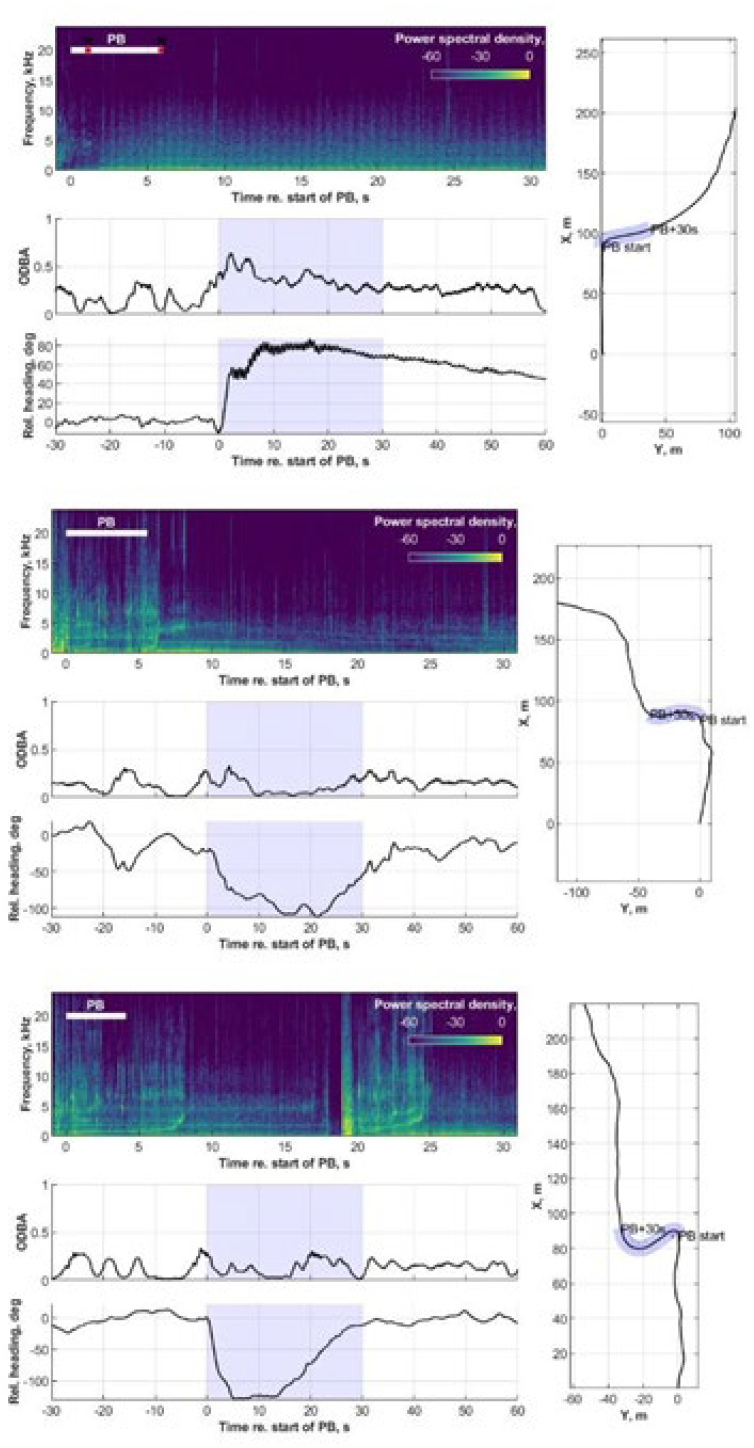
Three composite plots generated from DTAG data during playback, consisting of spectrogram (top), ODBA (middle), relative heading (bottom), and reconstructed movement track (right). Playback stimuli are difficult to discern in the spectrograms due both to their low RL and expanded time axis. However, a clear change in direction can be observed in both the relative heading and movement tracks. Note also that ODBA does not change noticeably in these trials. Top: adult male, who turned and swam away from a playback of NSWB; Middle: juvenile male, who turned and swam toward a playback of his own SW; Bottom: juvenile male who turned toward a playback of NSWA but remained in place

### Non-signature whistle playbacks

## Methods

As described above, the protocol for trials to free-swimming dolphins was to play back short sequences (consisting of two whistles separated by 2-3 sec) of different whistle types, each separated by at least 15 minutes of silence, to each target animal. Because our sample sizes of NSWA and NSWB were larger than other NSW types at this stage, we decided to emphasize exploring functions of these two whistle types. Our control stimuli were representative exemplars of the target animal’s own SW, to which we predicted dolphins would respond by approaching the speaker, as SW copies have been found to be an affiliative way of initiating contact with another dolphin (King & Janik, 2013; King et al. 2013). In randomized order, one whistle sequence consisted of two representative exemplars of the animal’s own SW (control), and the other was two exemplars of either NSWA or NSWB (randomly chosen from 8 exemplars of each whistle type, to avoid pseudoreplication). We also played back different exemplars of two other NSW types in 6 trials (shown in the 9^th^ and 10^th^ columns of Figure 8). As noted above, in instances where we stayed with a tagged animal for an extended period, we carried out a 3^rd^ (n=7) and 4^th^ trial (n=6); these stimuli consisted of an unfamiliar whistle or another NSW. Audio from the hydrophone, DTAG if used, and from a voice recorder of the primary observer were synchronized with drone video. Responses were scored as “positive” or “negative”, based on the animal’s initial orientation relative to the playback at its start. Trials in which responses were not observed or not clear were classified as “undetermined.”

## Results

In November 2023 and May 2024 we carried out 54 useable playback trials to 23 different target animals. Of these, 8 wore DTAGs in 24 trials. Details of these experiments are summarized in Table 1. Not all paired trials were completed, and some were discarded for a variety of reasons, resulting in uneven numbers of different types of trials.

**Table 1:**
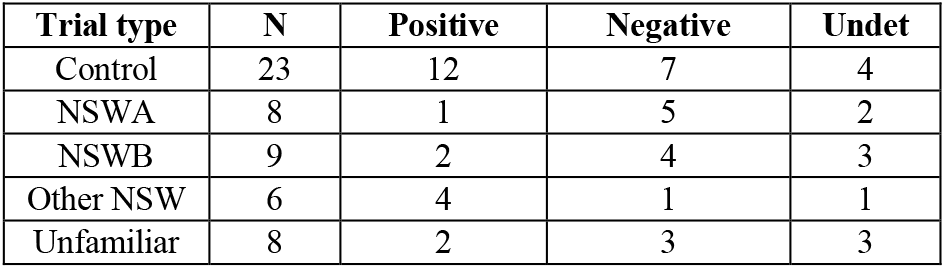
Summary of playback responses.

Control trials, in which we played the target dolphin’s own SW, resulted in positive responses in 63% of trials in which the response was determined. Playbacks of NSWA elicited negative responses in 83% of trials, whereas other NSW types elicited more positive responses (80% of trials where response was determined). NSWB and unfamiliar whistles elicited varying responses, with 3 of 4 positive responses being from males. In trials with tags, we were able to visualize changes in orientation (see examples in Figure 12), providing confirmation for observed responses in drone videos. However, ODBA was not found to significantly increase following these lower RL trials, providing further evidence for received level playing a large role in triggering some of the stronger responses that we observed in earlier trials.

## Discussion

This study provides the first evidence for widespread sharing of stereotyped NSWs in a dolphin communication system. NSWA seems likely to serve an alarm-type function, given that it elicited a majority of negative responses. Our suggested function of NSWB is a “query”-type whistle, produced when something unexpected or unfamiliar is heard. In the two instances where playbacks elicited production of NSWB, the two members of a male alliance were together at the time of playback, but heard copies of their own SW, which would normally be produced by their partner. With their partners by their sides, these copies of their own whistles were likely unexpected and perhaps difficult for the target animals to interpret. Similarly, the NSW playbacks to which we initially observed NSWB responses (Sayigh et al. 2017) were likely unexpected in the health-assessment context, in which dolphins typically produce approximately 85% SW (Sayigh et al. 2022). If our interpretation is correct, we would expect animals in different age and sex classes to use and respond to NSWB differently; for example, males may be more interested in investigating unfamiliar whistles in some contexts, whereas mothers and calves may avoid them. In fact, the two observed positive responses to NSWB playbacks were by males. Our playbacks also provide further support for the referential nature of SWCs, given that control SW stimuli elicited a majority of positive responses.

These preliminary results provide intriguing insights into the functions of different whistle types, however, they also highlight the complexity of the dolphin communication system, and thus the need for larger sample sizes before being able to draw definitive conclusions about functions of different NSW types. In reviewing drone videos, we have concluded that we should incorporate the additional response types “Interest” and “Ambivalent” to better describe some of the observed responses. For example, in some positive responses, animals turned toward the stimulus but then stayed in position, rather than approaching (e.g., Figure 12, bottom panel). In others, they initially approached but then turned away. These variations in responses highlight the need for more data to be able to correctly interpret them. In addition to these nuances in response types, we also found that individual dolphins may respond differently to the same whistle type, even copies of their own SW. This is not surprising - for example, not all humans respond identically to the same signal (even their name); rather responses depend greatly on individual characteristics (age, sex), context (behavioral state, companions or lack thereof, motivation to mate), and history of interactions (positive or negative) with other individuals. In our playbacks, one of the “ambivalent” responses involved a situation where the target animal was in a subgroup of 4 animals, with another subgroup of 4 nearby. Upon hearing the playback, the target animal’s subgroup turned and swam quickly toward the other subgroup. Thus, in a complex system such as this, we would not predict a “one-size-fits-all” response to every stimulus type, which makes it more difficult to study but also more interesting. However, we are optimistic that additional playback trials will shed light on the varied functions of shared NSWs in dolphins.

As mentioned above, bottlenose dolphins seem to use SW and SWC as learned referential labels for other individuals, but there is an outstanding question of whether referential communication is limited to other individuals. Given that dolphins are flexible, life-long vocal production learners, unlike most other non-human mammals, it is likely that shared, stereotyped NSW are both learned and referential, like SW and SWC. Our discovery and categorization of NSW in a wild population sets the stage for us to continue to test for contextual use of these shared signals, which will illuminate whether they are linked to other potential referents. Overall, our study provides the first evidence in dolphins for a wider repertoire of shared, context-specific signals, which could form the basis for a language-like communication system.

## Data availability

Sound and image files corresponding to the SW and NSW used as stimuli in the playback experiments described here are included in a data repository, with URI https://hdl.handle.net/1912/69819 and DOI 10.26025/1912/69819. Three examples each of 10 additional shared NSW are also included.

## Acknowledgments

Many people have contributed to this multidecadal effort, both in the field and in the laboratory. These include, but are not limited to, Blair Irvine, Michael Scott, and the staff, students, volunteers, and collaborators of the Sarasota Dolphin Research Program (SDRP), who made it possible to collect whistle data from, and provided long-term sighting, life history, and relationship data for the Sarasota Bay dolphins. SDRP was supported by Dolphin Quest, Inc., National Oceanographic and Atmospheric Administration, Disney, Harbor Branch Oceanographic Institute, The Royal Society, the Charles and Margery Barancik Foundation, and others. Multiple individuals at both the University of North Carolina Wilmington and Woods Hole Oceanographic Institution helped with field recordings and data extraction.

Financial support for the whistle database project has come from the Protect Wild Dolphins fund at Harbor Branch Oceanographic Institute, Vulcan Machine Learning Center for Impact, Allen Institute for Artificial Intelligence, Adelaide M. & Charles B. Link Foundation, and Dolphin Quest, Inc. Support for playback experiments was provided by the Davis Family and currently by the Avatar Alliance Foundation. Research was conducted under a series of Scientific Research Permits issued by the National Marine Fisheries Service, and was approved by Institutional Animal Care and Use Committees at Mote Marine Laboratory, University of North Carolina Wilmington, Hampshire College, and Woods Hole Oceanographic Institution.

## Footnotes

1 NSW playbacks were not included in this analysis because they were all carried out at lower source levels.

